# Fibroadipogenic Progenitor Cells Contribute to Tongue Skeletal Muscle Hypertrophy in Beckwith-Wiedemann Syndrome

**DOI:** 10.64898/2026.08.18.745577

**Authors:** Elisia D. Tichy, Shrey Pawar, Madison Newsome, Mara Fallon, Anna T. Nguyen, Gavriela Kalish-Schur, Mariah A. Byrne, Darryl Kinnear, Harry Kozakewich, Jennifer M. Kalish

## Abstract

Beckwith-Wiedemann syndrome (BWS) is a pediatric imprinting disorder characterized by tissue overgrowth, most commonly macroglossia, which can result in airway and feeding complications. Although dysregulated growth is a defining feature of BWS, the cellular interactions that drive organ-specific overgrowth remain poorly understood. We previously demonstrated that BWS macroglossia arises through distinct cell-intrinsic and cell-extrinsic mechanisms depending on molecular subtype. Here, we identify fibroadipogenic progenitor cells (FAPs) as modulators of myogenic differentiation and fusion in the human BWS tongue. BWS-derived FAPs were not increased in abundance *in situ* and did not exhibit hyperproliferation *in vitro*. Instead, FAPs from one BWS subtype promoted enhanced differentiation and fusion of normal human myoblasts. Secretome profiling revealed enrichment of CATHEPSIN L and TRANSFERRIN in conditioned media from these FAP populations, and functional perturbation of these factors supported their role in regulating myogenesis. These findings define a non-cell-autonomous mechanism of muscle overgrowth and implicate mesenchymal-myogenic signaling as a context-dependent driver of tissue expansion in an imprinting disorder.

## Introduction

Beckwith-Wiedemann syndrome (BWS) is a pediatric epigenetic overgrowth and cancer predisposition disorder caused by dysregulated imprinting at chromosome 11p15.5. This locus contains two imprinting control regions, imprinting center 1 (IC1) and imprinting center 2 (IC2), which exhibit parent-of-origin-specific methylation, with methylation on the paternal allele at IC1 and on the maternal allele at IC2 ^1^. These methylation marks regulate expression of growth-related genes within the IGF2/H19 and CDKN1C/KCNQ1OT1 domains, respectively, and disruption of this balance underlies the major molecular subtypes of BWS ^2^. Two of the most common molecular BWS subtypes are loss of methylation at imprinting center 2 (IC2 LOM) and paternal uniparental disomy of 11p15 (pUPD11). pUPD11 exhibits both IC2 LOM as well as IC1 gain of methylation (IC1 GOM) within the 11p15 region. These BWS-specific changes alter the expression of genes that regulate growth and development through distinct epigenetic mechanisms ^2,3^, yet how these changes are translated into tissue-specific overgrowth remains unclear.

Macroglossia, or tongue overgrowth, is one of the most common features of BWS, affecting approximately 80% of patients and often impairing breathing, feeding, speech, and sleep ^4^. Romeo *et al.* recently described an index of macroglossia severity for BWS, where it was found macroglossia severity varied by BWS subtype ^5^. Specifically, severe tongue overgrowth requiring reduction surgery was most frequently observed in the IC2 LOM subtype, followed by the pUPD11 subtype ^5^. IC1 GOM macroglossia rarely required tongue reduction surgery ^5^. In an effort to understand the drivers of BWS tongue overgrowth, a previous comparative study demonstrated that macroglossia in BWS is characterized by skeletal muscle hypertrophy in both IC2 LOM and pUPD11 subtypes, indicating convergent phenotypic outcomes ^6^. However, skeletal muscle progenitor cells isolated from IC2 LOM tongues, but not from pUPD11 tongues, intrinsically generated enlarged myotubes *in vitro*, even though both subtypes share disruption of the 11p15 IC2 region ^6^. This discrepancy suggests that, unlike IC2 LOM, pUPD11-associated tongue overgrowth may depend more strongly on extrinsic signals provided by the surrounding tissue microenvironment.

Skeletal muscle progenitors rely on paracrine cues from resident stromal populations to coordinate differentiation and fusion during muscle growth. Fibroadipogenic progenitor cells (FAPs) are mesenchymal lineage cells that influence myogenesis through their secretomes and thus represent a potential source of subtype-specific microenvironmental signals. Aberrant FAP behavior has been described in multiple skeletal muscle disorders, including fibrodysplasia ossificans progressiva ^7^ and muscular dystrophies ^8–11^, as well as in aging ^12,13^, where altered FAP properties contribute to impaired regeneration, fibrosis, adipogenesis, or loss of myogenic support. We therefore posited that pUPD11-associated macroglossia requires altered FAP-mediated cell-cell communication with myogenic cells, whereas IC2 LOM tongue overgrowth is less dependent on such paracrine regulation.

Here, we show that FAP abundance, collagen production, and proliferative capacity are comparable across control, IC2 LOM, and pUPD11 tongue tissues. FAP isolation and expansion *in vitro* revealed no hyperproliferative phenotype in BWS samples from either the IC2 LOM or pUPD11 subtype. To discern any BWS subtype-specific differences in FAP function, conditioned media (CM) was generated from primary FAPs cultured *in vitro* and applied to normal human primary myoblasts. We observed no differences in myoblast proliferation following exposure to control, IC2 LOM, or pUPD11 CM. However, CM from pUPD11 FAPs, but not from control or IC2 LOM FAPs, enhanced myogenic differentiation and fusion under differentiation conditions. Unbiased proteomic analysis identified increased secretion of CATHEPSIN L and TRANSFERRIN in pUPD11 FAPs, and functional perturbation of these factors supported their role in regulating myogenesis. These findings reveal a pUPD11 subtype-specific paracrine mechanism that promotes skeletal muscle hypertrophy in BWS macroglossia.

## Results

### FAP abundance and fibrotic features are not increased in BWS tongue tissue

To examine whether FAP abundance is altered in BWS macroglossia, we assessed expression of the canonical FAP marker PDGFRA^14^ in skeletal muscle regions of control, BWS IC2 LOM, and BWS pUPD11 tongue tissue (Fig. 1A). We found no significant differences in PDGFRA-positive cell abundance among groups (Fig. 1B), indicating that FAP populations are not expanded in BWS tongues at the time of resection. Because FAPs are major contributors to extracellular matrix deposition in skeletal muscle, we next assessed collagen content using picrosirius red staining (Fig. 1C). Fibrotic area was likewise comparable across control, IC2 LOM, and pUPD11 samples (Fig. 1D).

**Figure 1.**
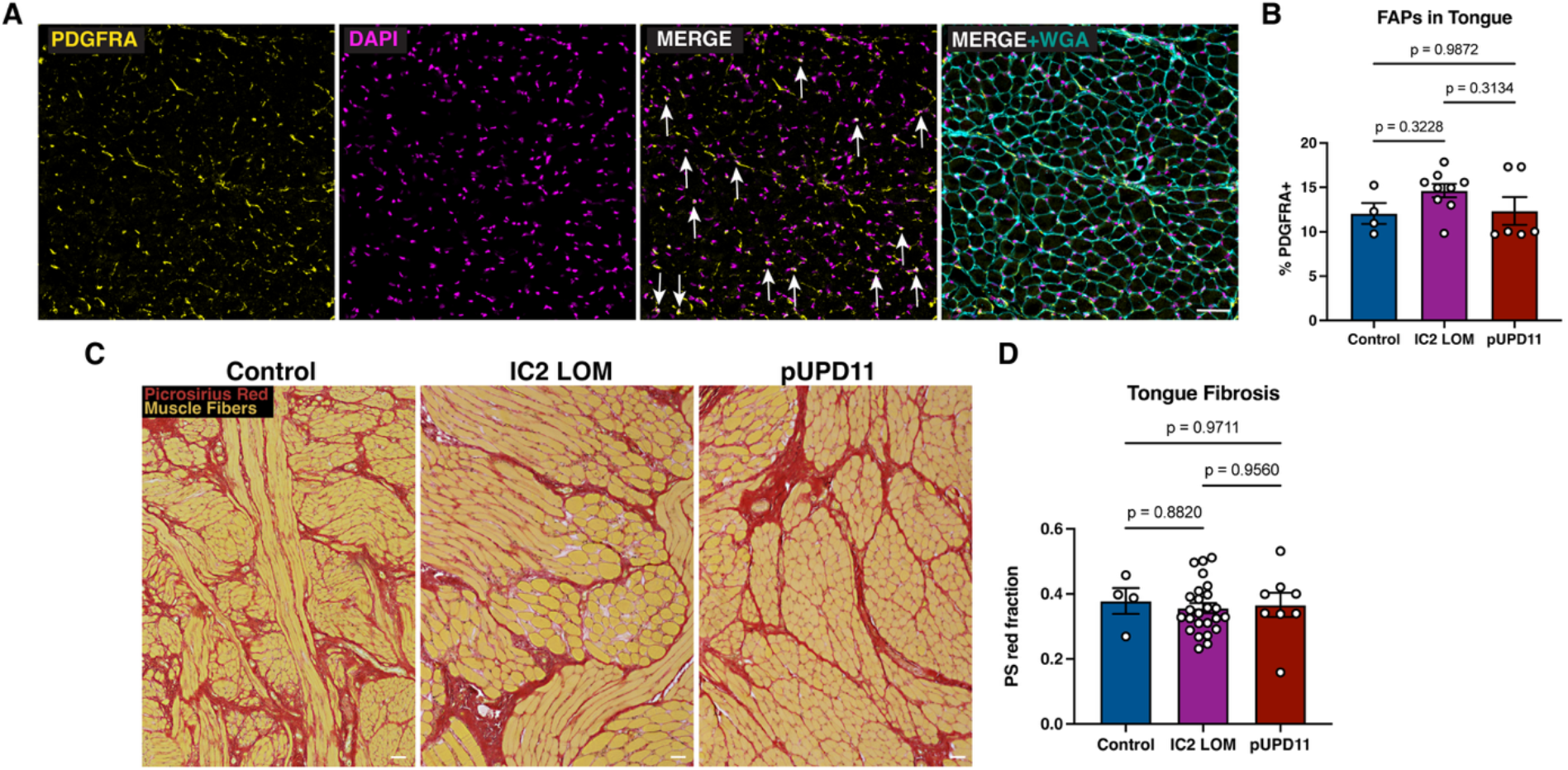
Characterization of FAPs in tongue tissue. **(A)** Representative cryosections of tongue tissue stained for PDGFRA (yellow) and nuclei (DAPI, magenta). Muscle was counterstained with WGA (cyan) to confirm proper FAP localization within skeletal muscle regions. Scale bar: 50 μm. **(B)** Quantification of PDGFRA-positive cells in tongue tissue. Each data point represents one unique patient sample. **(C)** Representative images of picrosirius red staining of FFPE tongue sections. Scale bar: 50 μm. **(D)** Quantification of fibrosis in tongue tissue. Each data point represents one unique patient sample. Statistical significance was determined by ANOVA.

### BWS tongue-derived FAPs retain canonical identity and do not exhibit enhanced proliferation

We next isolated live FAPs from cryobanked control, IC2 LOM, and pUPD11 tongue specimens by magnetic enrichment for PDGFRA. Purified cells expressed PDGFRA protein (Fig. S1A) and showed comparable expression of established FAP markers, including *PDGFRA*, *DCN*, *OSR1*, *COL1A1*, and *NT5E*, by qPCR (Fig. S1B-F). To confirm lineage identity, we assayed adipogenic potential. Following adipogenic induction, isolated cells accumulated neutral lipids detectable by BODIPY staining (Fig. S2A) and expressed adipogenic markers *LEP*, *ADIPOQ*, and *CEBPB* (Fig. S2B-D). These results support successful isolation of tongue-derived FAPs from all groups.

Because BWS is an overgrowth disorder, we next tested whether BWS-derived FAPs exhibit enhanced proliferation *in vitro*. EdU incorporation assays revealed no significant differences between control, IC2 LOM, and pUPD11 FAPs at a 2 h labeling interval (Fig. 2A-B). Cell cycle profiling similarly showed no increase in S-phase populations in BWS FAPs relative to controls (Fig. 2C-D).

**Figure 2.**
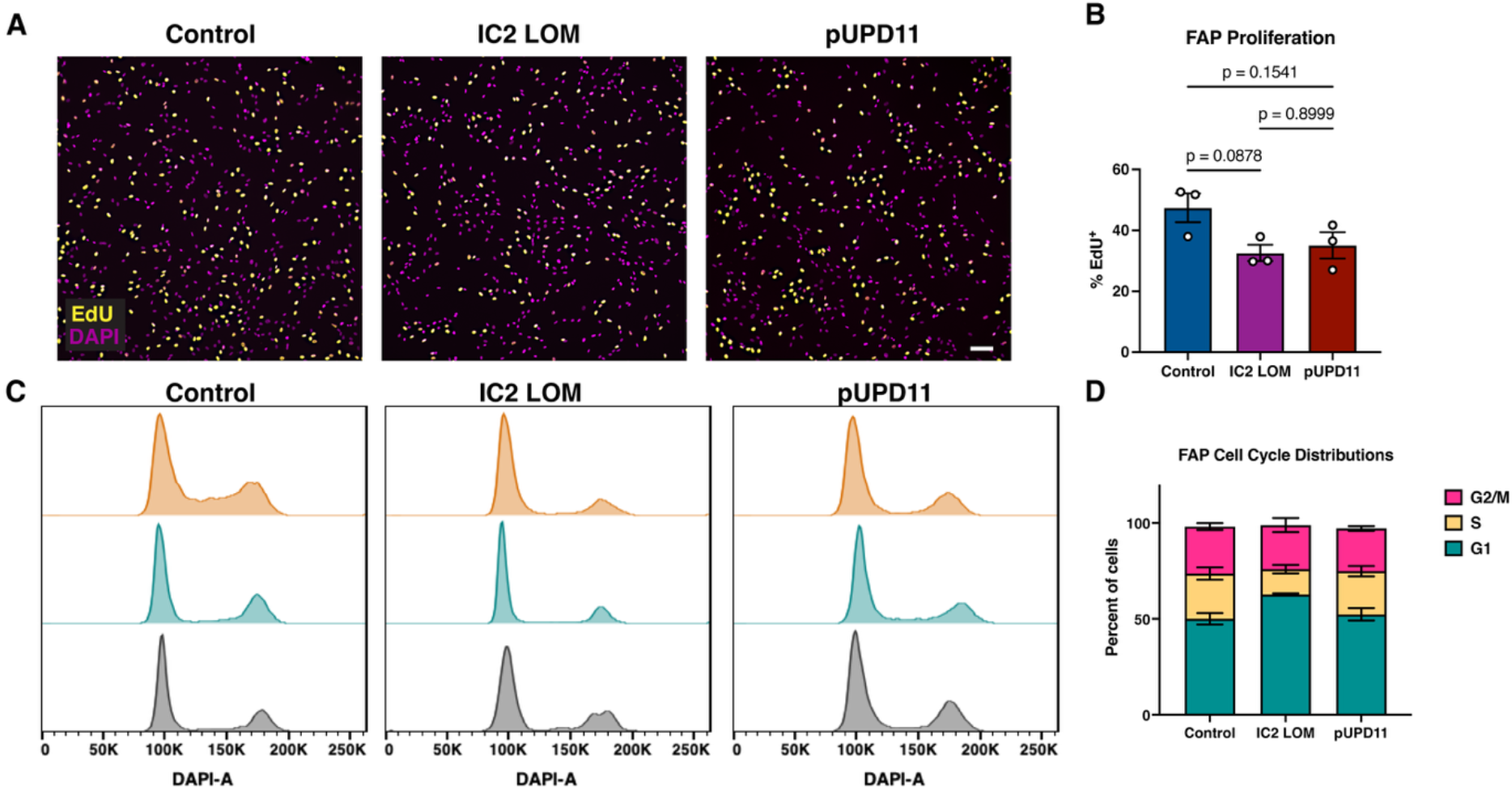
Tongue-derived FAPs do not exhibit enhanced proliferation in vitro. **(A)** Representative images of tongue-derived FAPs following EdU incorporation. Scale bar: 100 μm. **(B)** Quantification of FAP proliferation. n = 3 unique patient samples per group. **(C)** Cell-cycle profiles of asynchronously growing FAPs from control, IC2 LOM, and pUPD11 samples. Each graph is derived from one unique patient. **(D)** Quantification of cell-cycle distribution shown in (C).

### pUPD11 FAP-conditioned media does not alter myoblast proliferation

FAPs influence myogenic progenitor behavior through paracrine and microenvironmental signaling, with reported effects on proliferation, regenerative support, differentiation, and fusion ^15^. To determine whether FAP-derived secreted signals differ across groups, we generated CM from control, IC2 LOM, and pUPD11 FAPs and applied these to two primary human myoblast lines, one male and one female (Table S1). Effects of FAP CM on myoblast proliferation were assessed by EdU incorporation (Fig. 3A). We observed no significant differences in myoblast proliferation following exposure to CM from control, IC2 LOM, or pUPD11 FAPs in either myoblast line (Fig. 3B).

**Figure 3.**
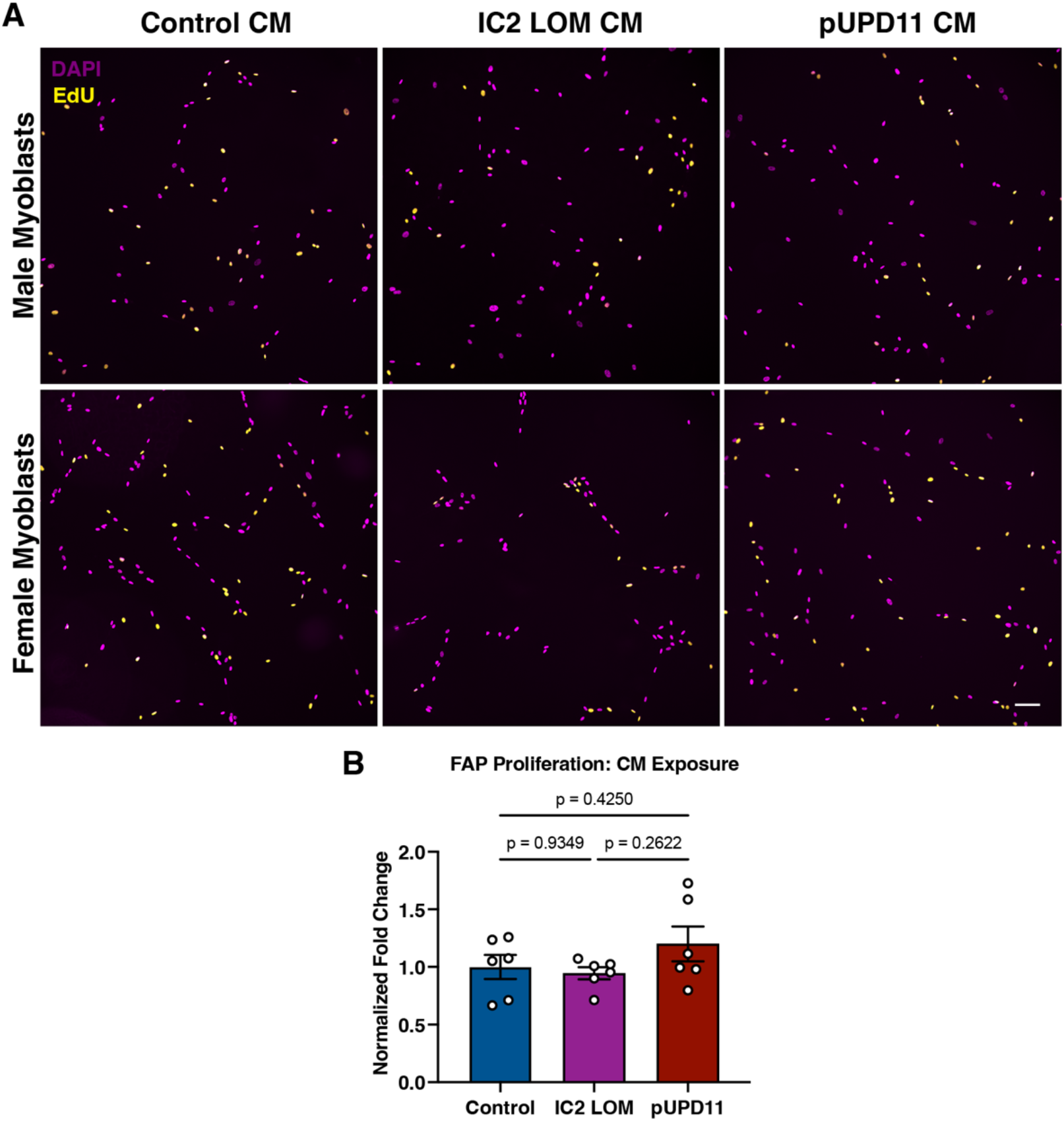
Effects of FAP CM on myoblast proliferation. **(A)** Representative images of primary human myoblasts, one male line and one female line, exposed to CM from control, IC2 LOM, or pUPD11 FAPs and assayed for proliferation by EdU incorporation. Scale bar: 100 μm. **(B)** Quantification of EdU incorporation in male and female myoblast lines. Each data point represents one combination of myoblast line and FAP CM source. Data were normalized to the control condition for each respective myoblast donor sex. Statistical significance was determined by ANOVA.

### pUPD11 FAP-conditioned media enhances myogenic differentiation and fusion

Using the same primary human myoblast system, we then assessed differentiation and fusion post-FAP CM exposure using MYOGENIN and MYOSIN HEAVY CHAIN 4 as markers of early and late myogenesis, respectively (Fig. 4A). The proportion of MYOGENIN-positive nuclei did not differ significantly between myoblasts exposed to control and IC2 LOM FAP CM (Fig. 4B). Myoblasts exposed to pUPD11 FAP CM showed significantly increased MYOGENIN positivity relative to controls and a similar upward trend relative to IC2 LOM samples (Fig. 4B). Fusion indices were significantly increased in pUPD11 CM relative to both control and IC2 LOM CM, whereas no differences were observed between control and IC2 LOM FAP CM-exposed cells (Fig. 4C).

**Figure 4.**
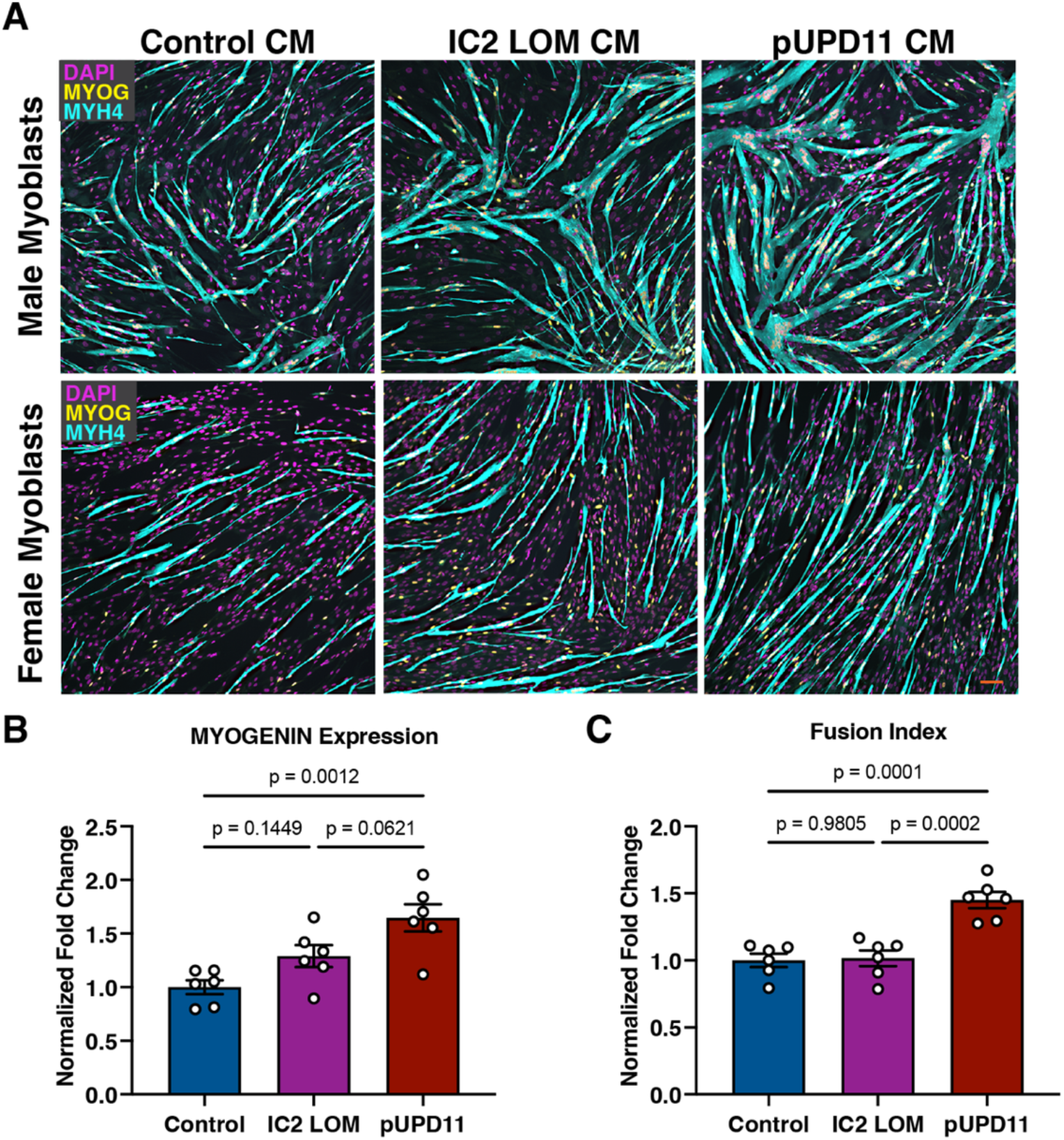
Effects of FAP CM on myoblast differentiation and fusion. **(A)** Representative images of primary human myoblasts, one male line and one female line, exposed to CM from control, IC2 LOM, or pUPD11 FAPs and assayed for differentiation by MYOGENIN staining and for fusion index. Scale bar: 100 μm. **(B)** Quantification of MYOGENIN-positive nuclei in normal male and female myoblast lines. Each data point represents one combination of myoblast line and FAP CM source. Data were normalized to the control condition for each respective myoblast donor sex. **(C)** Quantification of fusion index in FAP CM-exposed myoblasts. Statistical significance was determined by ANOVA.

### Secretome profiling identifies candidate pro-myogenic factors enriched in pUPD11 FAPs

To identify candidate mediators of the effect of pUPD11 FAP CM on myoblast differentiation and/or fusion, we performed unbiased mass spectrometry on FAP CM from all groups. We found similar total protein recovery across samples, which argued against major collection bias (Fig. S3A). Principal component analysis showed that control and IC2 LOM samples clustered relatively closely, whereas pUPD11 samples were more distinct (Fig. S3B).

Across comparisons, relatively few proteins differed between control and IC2 LOM secretomes, whereas pUPD11 samples displayed broader divergence from both other groups (Fig. S4A). Gene ontology cellular localization analysis showed broadly similar distributions of proteins across experimental groups, including extracellular, plasma membrane, cytoplasmic, and nuclear categories (Fig. S4B-S4D). Because our goal was to identify secreted mediators of cell-cell communication, we therefore focused subsequent analyses on proteins annotated to extracellular or plasma membrane compartments.

Within this filtered set, comparison of IC2 LOM and control secretomes yielded only a small number of significant candidates (Fig. 5A). Downregulated proteins in IC2 LOM included DKK3 and RNASET2. DKK3 has been linked to skeletal muscle atrophy when overexpressed, both *in vitro* and *in vivo* ^16^, whereas RNASET2 has recently been implicated as a macrophage-derived factor that promotes muscle stem cell fusion ^17^.

**Figure 5.**
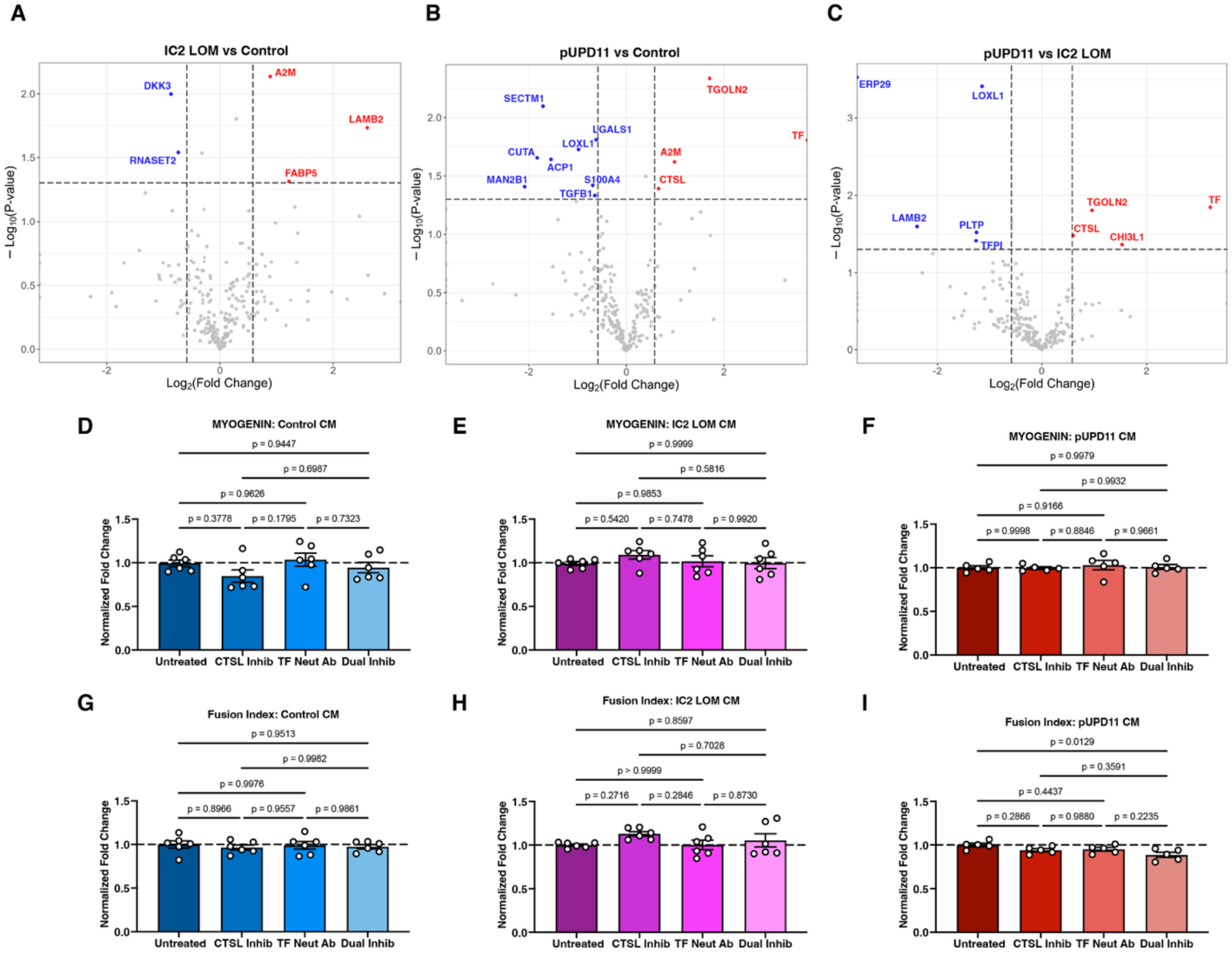
Identification and functional testing of putative pUPD11 FAP-secreted regulators of late stage myogenesis. **(A)** Volcano plot comparing mass spectrometry data from IC2 LOM and control FAP secretomes. **(B)** Volcano plot comparing mass spectrometry data from pUPD11 and control FAP secretomes. **(C)** Volcano plot comparing mass spectrometry data from pUPD11 and IC2 LOM FAP secretomes. **(D-F)** Quantification of MYOGENIN-positive nuclei in normal male and female human myoblast lines following exposure to FAP-conditioned differentiation medium from control, IC2 LOM, or pUPD11 FAPs, respectively. Inhibitors, alone or in combination, were added at the time of differentiation onset. Each data point represents one biological replicate. Note: High-dose CATHEPSIN L inhibitor was used in myoblast-only loss-of-function assays; a 100-fold lower dose was used in FAP-conditioned media experiments to minimize direct effects on intrinsic myoblast differentiation. Data were normalized to the untreated samples from each condition. Trendline was set to 1. **(G-I)** Quantification of myogenic fusion in normal male and female human myoblast lines following exposure to FAP-conditioned differentiation medium from control, IC2 LOM, or pUPD11 FAPs, respectively. Inhibitors, alone or in combination, were added at the time of differentiation onset. Each data point represents one biological replicate. Data were normalized to the untreated samples from each condition. Trendline was set to 1.

Upregulated proteins in IC2 LOM included A2M, LAMB2, and FABP5. A2M is a broad extracellular protease regulator and has been reported to increase during early differentiation of human myoblasts derived from aged individuals ^18^. LAMB2 contributes to neuromuscular junction organization ^19^, whereas FABP5 is linked to lipid handling, although its direct role in skeletal muscle differentiation or fusion remains unclear ^20^. Because CM from IC2 LOM FAPs produced only modest differences in myogenic output compared to control CM, we did not pursue these candidates further.

In contrast, pUPD11 secretomes showed broader changes relative to both control and IC2 LOM samples. Compared with control, pUPD11 CM contained reduced levels of SECTM1, CUTA, MAN2B1, ACP1, LOXL1, LGALS1, S100A4, and TGFB1, and increased levels of CTSL, A2M, TGOLN2, and TF (Fig. 5B). Direct comparison of pUPD11 and IC2 LOM similarly identified increased TGOLN2, TF, CHI3L1, and CTSL in pUPD11 secretomes and reduced ERP29, LOXL1, LAMB2, PLTP, and TFPI (Fig. 5C).

Several of the proteins reduced in pUPD11 secretomes have been linked to skeletal muscle remodeling. For example, LOXL1, LGALS1, and S100A4 have been identified in proteomic studies of skeletal muscle remodeling after exercise ^21^. Notably, TGFB1 levels were slightly reduced in pUPD11 CM relative to control samples. TGFB signaling is a well-established inhibitor of skeletal muscle differentiation and myoblast fusion ^22^. However, because TGFB1 was not differentially reduced between pUPD11 and IC2 LOM samples, decreased TGFB1 alone is less likely to account for the pUPD11-specific enhancement of myogenesis.

Because treatment with pUPD11 CM led to significant changes in myoblast differentiation and fusion, we next aimed to functionally validate pUPD11-enriched secreted proteins *in vitro* through recombinant protein addition or inhibition strategies during normal myogenesis. Among these proteins, TGOLN2 and CHI3L1 were of potential interest but have less direct links to myogenic differentiation. TGOLN2 is a Golgi-associated cargo-binding protein without a defined role in skeletal muscle differentiation. CHI3L1 has been shown to promote proliferation of myogenic cells during exercise responses but does not appear to regulate expression of myosin heavy chain during differentiation ^23^. By contrast, TRANSFERRIN (TF) and CATHEPSIN L (CTSL) were enriched in pUPD11 secretomes relative to both comparison groups and have stronger biological precedent for regulating muscle growth or differentiation.

TRANSFERRIN has long been recognized as a positive regulator of skeletal muscle differentiation ^24^. Genetic disruption of transferrin receptor 1 in skeletal muscle of rodents reduces muscle volume and myofiber size and leads to postnatal failure to thrive ^25^. Satellite cell-specific deletion of transferrin receptor 1 impairs proliferation and fusion during regeneration ^26^. CATHEPSIN L is a lysosomal cysteine protease that can also be secreted into the extracellular environment and participate in matrix remodeling ^27^. In skeletal muscle, Cathepsin L expression increases during mouse myoblast differentiation^28^ and altered CATHEPSIN L activity has been associated with multiple muscle remodeling states including diabetes, fasting, and muscular dystrophy ^29–33^. Together, these observations supported prioritization of TRANSFERRIN and CATHEPSIN L as candidate pUPD11 FAP-derived regulators of myogenesis.

### CATHEPSIN L and TRANSFERRIN contribute to human myogenic differentiation and fusion

To determine whether candidate factors enriched in pUPD11 FAP secretomes directly influence myogenesis, we performed gain-of-function experiments in primary human myoblast cultures. Recombinant active CATHEPSIN L or holo-TRANSFERRIN was added during myoblast differentiation. Addition of active CATHEPSIN L had no detectable effect on MYOGENIN expression or fusion (Fig. S5A, S5B). By contrast, addition of holo-TRANSFERRIN significantly increased both early and late myogenesis in both normal primary myoblast lines (Fig. S5C, S5D).

We next asked whether endogenous CATHEPSIN L and TRANSFERRIN activity is required for myogenic differentiation by performing loss-of-function experiments in normal primary human myoblasts under standard differentiation conditions. Pharmacologic inhibition of CATHEPSIN L using a high dose of the irreversible inhibitor Z-FY(t-Bu)-DMK significantly reduced MYOGENIN expression and fusion index (Fig. S5E, S5F), indicating that CATHEPSIN L activity contributes to myogenic progression *in vitro*. By contrast, neutralization of TRANSFERRIN did not alter myoblast differentiation or fusion under standard differentiation conditions (Fig. S5G, S5H). Because high-dose CATHEPSIN L inhibition directly suppressed myogenesis in myoblast-only assays, we used a 100-fold lower concentration of inhibitor in subsequent conditioned-media experiments to minimize direct effects on intrinsic myoblast differentiation and instead test whether CATHEPSIN L-dependent activity contributes to the pro-myogenic effects of FAP-conditioned media. At this lower concentration, CATHEPSIN L inhibition alone, TRANSFERRIN inhibition alone, or dual inhibition had no significant effect on MYOGENIN expression across control, IC2 LOM, or pUPD11 CM conditions (Fig. 5D-5F). Fusion indices were likewise unchanged in control and IC2 LOM CM conditions (Fig. 5G, 5H). In contrast, although single inhibition of CATHEPSIN L or TRANSFERRIN did not significantly reduce fusion in pUPD11 CM-treated myoblasts, combined inhibition produced a slight but significant reduction in fusion index relative to untreated pUPD11 CM samples (Fig. 5I). These data suggest that CATHEPSIN L and TRANSFERRIN act as complementary contributors to the pro-fusion activity of pUPD11 FAP-conditioned media.

## Discussion

In this study, we identify a subtype-specific stromal secretory mechanism that contributes to macroglossia in BWS. Although we previously found that both IC2 LOM and pUPD11 tongues exhibit skeletal muscle hypertrophy, our data support the idea that pUPD11-associated overgrowth is shaped in part by the local tissue microenvironment. Specifically, we show that FAP abundance and fibrosis are not increased in BWS tongues, but that CM from pUPD11 FAPs enhances myogenic differentiation and fusion of normal human myoblasts. These findings support a model in which altered FAP function, rather than overt expansion of the FAP compartment, contributes to skeletal muscle hypertrophy in this BWS subtype.

The absence of increased FAP abundance or proliferation suggests that altered FAP secretory behavior, rather than intrinsic growth differences in FAPs, underlies the phenotype. This distinction is important because previous work in BWS tongue has suggested that IC2 LOM and pUPD11 subtypes reach a similar hypertrophic endpoint through different mechanisms ^6^. Our findings are consistent with that model and extend it by implicating pUPD11 FAPs as paracrine regulators of skeletal muscle growth. In this framework, altered stromal-myogenic communication within the tongue microenvironment amplifies myogenic differentiation and fusion and thereby contributes to tissue overgrowth.

This interpretation is also consistent with a growing body of omics work showing that FAPs undergo context-dependent changes in gene expression and signaling across regeneration, aging, and disease, and that these changes are likely to influence how FAPs communicate with neighboring myogenic cells ^8,34–38^. Bulk transcriptomic and proteomic studies in murine muscle have identified marked changes in FAP-associated pathways during acute injury, dystrophy, and metabolic perturbation ^8,35,36^, while single-cell and single-nucleus transcriptomic studies in mouse and human skeletal muscle have further highlighted substantial FAP heterogeneity and predicted secretory programs associated with regeneration or pathology ^34,37,38^. In human disease settings, omics-guided analyses in Duchenne muscular dystrophy have begun to define disease-associated signaling pathways in FAPs, and recent engineered human skeletal muscle studies show that FAPs can enhance structural and functional muscle properties, supporting the idea that FAPs influence myogenic outcomes through niche-associated and paracrine mechanisms ^39,40^. Notably, direct proteomic studies of FAPs remain relatively limited. This likely reflects, at least in part, the technical difficulty of isolating and expanding sufficient FAPs for protein-level analyses, particularly from limited human skeletal muscle tissue. In this context, our direct proteomic analysis of BWS tongue FAP-conditioned media provides a complementary biochemical view of disease-associated FAP signaling and identifies candidate secreted factors linked to subtype-specific myogenic outcomes. To our knowledge, CATHEPSIN L and TRANSFERRIN have not emerged as established FAP-derived regulators of myogenesis in prior FAP omics studies, supporting the novelty of these candidates in the BWS tongue microenvironment.

Proteomic analysis further supported this model by identifying broader secretome remodeling in pUPD11 FAPs than in IC2 LOM FAPs. Although only a small number of extracellular or plasma membrane-associated proteins differed between control and IC2 LOM secretomes, pUPD11 FAP secretomes showed broader divergence from both other groups. Reduced TGFB1 in pUPD11 CM is consistent with a more permissive fusion environment, given the established role of TGFB signaling as a repressor of myogenic differentiation and fusion^22^. However, because TGFB1 was not differentially reduced between pUPD11 and IC2 LOM samples, it is unlikely to fully explain the pUPD11-specific phenotype. Among proteins increased in pUPD11 CM, TRANSFERRIN and CATHEPSIN L emerged as the strongest candidates on the basis of both differential abundance and prior links to muscle biology.

The combined inhibition data further support the idea that TRANSFERRIN and CATHEPSIN L act as complementary components of the pUPD11 FAP secretome rather than as fully redundant regulators. Although inhibition of either factor alone had only modest effects in the context of pUPD11 CM, combined inhibition slightly but significantly reduced myogenic fusion, consistent with parallel contributions to the pro-myogenic phenotype.

TRANSFERRIN has well-established roles in skeletal muscle differentiation^24^, and our gain-of-function experiments are consistent with that literature. Addition of holo-TRANSFERRIN enhanced both MYOGENIN expression and myotube fusion, supporting the interpretation that increased TRANSFERRIN in the pUPD11 FAP secretome contributes to the pro-myogenic phenotype. By contrast, TRANSFERRIN neutralization did not alter myogenesis under standard differentiation conditions, indicating that a requirement for endogenous TRANSFERRIN was not detectable in this assay. This may reflect ligand abundance, iron availability, neutralization efficiency, receptor occupancy, or compensatory pathways under baseline culture conditions.

CATHEPSIN L showed a different pattern. Exogenous active CATHEPSIN L did not significantly enhance differentiation or fusion, whereas high-dose pharmacologic inhibition of CATHEPSIN L reduced both MYOGENIN expression and myotube formation. This apparent asymmetry suggests that CATHEPSIN L may function as a context-dependent regulator of myogenesis rather than as a simple dose-dependent enhancer. One possible explanation is that CATHEPSIN L exhibits optimal activity at acidic pH and reduced activity under neutral conditions, consistent with its canonical lysosomal role, while extracellular activity may depend on local binding partners, matrix context, or other stabilizing microenvironmental factors ^41^. In that setting, inhibition would suppress endogenous CATHEPSIN L activity that is already present or locally stabilized during differentiation, whereas simple addition of recombinant enzyme to culture medium may not recapitulate its physiological mode of action. Thus, our data support a role for CATHEPSIN L in late myogenesis by acting on the extracellular environment. In addition, these findings raise the possibility that CATHEPSIN L represents a previously underappreciated regulator of late-stage myogenesis.

To more closely examine the relationship of CATHEPSIN L and TRANSFERRIN in the context of pUPD11 FAP secretomes, we carried out inhibition experiments, alone and in combination, using FAP CM. In this context, we lowered the concentration of CATHEPSIN L inhibitor to have less direct effects on intrinsic myogenic properties of normal human myoblasts, and instead preferentially target these factors in the CM. We found a significant reduction in fusion index when myoblasts were treated with pUPD11 CM together with the combination of inhibitors. This result was not observed in control or IC2 LOM samples treated under the same conditions, supporting a role for TRANSFERRIN and CATHEPSIN L in the context of pUPD11-driven macroglossia.

More broadly, these findings provide a plausible explanation for why pUPD11 satellite cells do not exhibit overt intrinsic abnormalities despite the presence of macroglossia in patients. Rather than arising from a strong cell-autonomous defect in the myogenic progenitor population, as in IC2 LOM ^6^, pUPD11-associated myogenic tongue overgrowth may be driven primarily by extrinsic cues from FAPs.

This study has several limitations. The sample size is small because of the rarity of primary human BWS tongue specimens. In addition, our *in vitro* CM assays were designed to isolate stromal-myogenic signaling, but they do not fully recapitulate the cellular and extracellular complexity of the native tongue microenvironment. Our proteomic prioritization strategy also focused on the subset of differentially abundant extracellular factors, and additional secreted components, such as extracellular vesicles containing microRNAs ^42,43^, may contribute to the pro-myogenic phenotype. Finally, because the analyzed tissues were obtained after macroglossia was already established, we cannot determine whether the same signaling interactions operate during the earlier developmental stages when tongue overgrowth is initiated.

In summary, our data support a model in which pUPD11-associated macroglossia is driven in part by altered stromal-myogenic communication within the tongue microenvironment. Although FAPs are neither expanded nor intrinsically hyperproliferative in BWS tongues, a subset of FAPs derived from pUPD11 tissues secretes factors that enhance myoblast differentiation and fusion, thereby promoting skeletal muscle hypertrophy. These findings define a non-cell-autonomous mechanism of muscle overgrowth and highlight FAP-derived signaling as a regulator of organ-specific expansion in an imprinting disorder.

## Materials and Methods

### Patient samples

Human tongue specimens from patients with molecularly confirmed BWS were obtained from the BWS Registry and BioRepository at the Children’s Hospital of Philadelphia under Institutional Review Board approval (#13-010658) following informed consent. Tongue tissue was collected at the time of clinically indicated tongue reduction surgery. BWS subtypes included IC2 LOM and pUPD11. Methylation percentages at IC1 and IC2 for BWS tongue specimens used to generate FAPs were determined by the University of Pennsylvania Genetic Diagnostic Laboratory (Table S1).

Nonsyndromic/control pediatric autopsy tongue samples were obtained under institutional policies governing research use of de-identified specimens and were deemed exempt from IRB oversight. Patient demographic and molecular characteristics are provided in Fig. S6. Unless otherwise stated, biological replicates represent independent patient specimens.

### Tissue processing

Tongue tissue was processed by formalin fixation and paraffin embedding (FFPE), embedded in optimal cutting temperature compound (OCT) for cryosectioning, or cryopreserved in CryoStor CS10 (BioLife Solutions 210502) for *in vitro* assays. FFPE sections were prepared by the CHOP pathology core. Cryosections were cut at 10 μm and stored at -80°C until use. Tissue designated for cell isolation was cryopreserved in liquid nitrogen and thawed immediately prior to enzymatic digestion.

### Immunostaining of FAPs in human tongue tissue

Cryosections from control, IC2 LOM, and pUPD11 specimens (n = 3 per group) were fixed in 4% paraformaldehyde for 10 min, permeabilized with 0.5% Triton X-100 in PBS, and blocked in 3% BSA with 0.1% Triton X-100. Sections were incubated overnight at 4°C with goat anti-human PDGFRA (R&D Systems AF-307-NA, 1:100), followed by donkey anti-goat Alexa Fluor 555 (ThermoFisher A21432, 1:300) secondary antibody staining. Wheat germ agglutinin-Alexa Fluor 647 (Fisher W32466, 1:1000) was used to delineate myofiber boundaries. Nuclei were counterstained with DAPI. Images were acquired using a Nikon Ti2 widefield microscope with identical acquisition settings across samples. For each specimen, at least three nonoverlapping fields from three independent sections were analyzed. PDGFRA+ cells were manually quantified using FIJI. Quantification was performed blinded to genotype.

### Tongue picrosirius red staining and analysis

Six-micron FFPE sections from control and BWS tongues were stained with picrosirius red (PSR) to assess collagen deposition. Images were acquired using an EVOS microscope under identical exposure settings. At least three fields per section and three sections per specimen were analyzed. Fibrotic area was quantified using the PSR_quantify ImageJ macro as previously described ^20^. Fibrosis fraction was calculated as the percentage of PSR-positive area relative to total tissue area. Quantification was performed blinded to genotype.

### FAP isolation and culture

FAPs were isolated from cryopreserved tongue tissue (n = 3 independent patients per group) using enzymatic digestion followed by magnetic-activated cell sorting (MACS). Tissue was digested with Collagenase IA (Sigma C9891) and Dispase II (Millipore D4693), mechanically dissociated, filtered through a 40 μm strainer, and subjected to red blood cell lysis (Sigma R7757).

Cells were incubated with goat anti-human PDGFRA antibody (R&D Systems BAF322) followed by anti-goat magnetic microbeads (Rockland) and purified using LS columns (Miltenyi Biotec). PDGFRA+ cells were eluted and plated in FAP growth media comprised of DMEM/F12 supplemented with 20% fetal bovine serum, 1X GlutaMAX, 1X antibiotic-antimycotic, and Primocin (Invivogen ant-pm-1). FAPs were used at passages 3-5 for all experiments.

### FAP immunostaining *in vitro*

Twenty thousand FAPs were plated on 8-well chamber slides and grown for 24 h in FAP proliferation medium. Cells were fixed in 4% paraformaldehyde in PBS for 5 min, permeabilized with 0.5% Triton X-100 in PBS for 5 min, blocked for 1 h at room temperature, and incubated overnight at 4°C with goat anti-human PDGFRA antibody (R&D Systems AF-307-NA). Cells were washed with PBS and incubated in buffer containing donkey anti-goat Alexa Fluor 647-conjugated secondary antibody (1/1000, ThermoFisher A21447) for 1 h at room temperature in the dark. Cells were washed with PBS, chambers removed, and coverglass was mounted using Prolong gold with DAPI (ThermoFisher P36935). Slides were imaged with a Nikon Eclipse TI32 fluorescence microscope. Representative images were taken to demonstrate enrichment of PDGFRA+ cells (FAPs).

### Cell cycle analysis

Asynchronously growing FAPs were harvested at 50% confluence. Briefly, cells were washed with PBS, detached with TrypLE Express, and centrifuged at 300 rcf for 5 min. The pellet was washed with PBS and fixed with ice-cold 70% ethanol overnight at 4°C. Cells were centrifuged at 600 rcf for 5 min, washed with 1% BSA in PBS, and stained with 1 μg/mL DAPI. Cells were analyzed for cell cycle position on a BD Fortessa with a violet laser and a 450/50 bandpass filter. Cell cycle analysis was conducted in FlowJo v.10.10.0.

### FAP adipogenesis

Confluent FAPs were differentiated to adipocytes using the adipogenic differentiation media kit (Lonza PT-3004), using three rounds of induction and maintenance, according to the manufacturer’s instructions. To confirm adipogenesis, undifferentiated or differentiated FAPs were incubated with 2.5 μM BODIPY 493 (Invitrogen D3922) in PBS for 30 min at 37°C. Nuclei were stained with Hoechst 33342 (5 ng/mL) and imaged on an EVOS M7000 microscope.

### FAP quantitative PCR

RNA was isolated from FAPs using Trizol and the Monarch Spin RNA Cleanup Kit (NEB T2040L). cDNA was generated using the Protoscript II cDNA synthesis kit (NEB E6300S). Quantitative real-time PCR was conducted on a QuantStudio 6 Pro using Taqman 2X Universal PCR master mix (ThermoFisher 4304437), in 20 μL reactions. Primers were all from ThermoFisher and included: *PDGFRA*-FAM (Hs00998018_m1), *COL1A1*-FAM (Hs00164004_m1), *DCN*-FAM (Hs00754870_s1), *OSR1*-FAM (Hs01586544_m1), *NT5E*(CD73)-FAM (Hs00159686_m1), *CEBPB*-FAM (Hs00942496_s1), *LEP*-FAM (Hs00174877_m1), *ADIPOQ*-FAM (Hs00977214_m1), and *UBC*-VIC; primer limited (Hs05002522_g1). Relative expression was determined by the ddCT method. For negative control for some assays, cDNA from HEK cells was used to set baseline.

### FAP proliferation assessments

Twenty thousand FAPs were plated in 8-well chamber slides in FAP proliferation media. After 24 h, 10 μM EdU was added to the media and incubated at 37°C for 2 h. Media was aspirated and cells were fixed in 4% paraformaldehyde in PBS for 5 min. Cells were washed with PBS and permeabilized with 0.5% Triton X-100 in PBS, followed by staining with the Click-iT EdU Alexa Fluor 488 Imaging kit (Fisher C10637), according to the manufacturer’s instructions. Chambers were removed from the slides and coverglass was mounted with Prolong gold with DAPI (ThermoFisher P36935). Slides were imaged on a Nikon Ti2 widefield fluorescent microscope.

### Myoblast culture and FAP CM exposure

Primary human myoblasts were purchased from Lonza (CC-2580). Donor information is provided in Supplementary Table 1. Myoblasts were expanded in SkGM-2 growth media (Lonza CC-3245) to passage 3 or passage 4. To generate FAP-CM, 1×10^6^ FAPs were plated onto 10-cm tissue culture dishes and grown in either FAP proliferation medium or myoblast differentiation medium (DMEM/F12, 5% horse serum, 1X Glutamax, 1X anti-anti, 1X primocin) for 48 h. FAP-CM was aliquoted and stored at -80°C until use.

To assay the effects of FAP-secreted factors on myoblast proliferation, 10,000 myoblasts were plated per well in 8-well chamber slides (Millipore-Sigma PEZGS0816). After 24 h, the media was changed to FAP-conditioned proliferation media and cells were cultured for another 48 h. EdU was added at a final concentration of 10 μM and was allowed to incorporate for 2 h. EdU incorporation was assessed as described above.

To query FAP-secreted factors on myoblast differentiation, differentiation media exposed to FAPs was applied to myoblasts plated in 8-well chambers slides (80,000 cells/well). After 3 days exposure, myoblasts were fixed in 4% paraformaldehyde in PBS for 5 min and immunostained as above using antibodies against MYOGENIN (abcam ab124800 1/100) and MYOSIN HEAVY CHAIN 4 (Fisher 12-6503-82); 1/100. Differentiation potential was defined as the percent of nuclei that were MYOGENIN positive. Fusion index was defined as the number of nuclei present in myosin heavy chain 4 positive myotubes (containing at least 2 nuclei) divided by the total numbers of nuclei.

### Mass spectrometry of tongue FAP-secreted factors

FAPs from 11 human tongue specimens, including four control, three IC2 LOM, and four pUPD11 samples, were plated at 1×10^6^ cells per 10-cm plate in FAP proliferation medium overnight. The following day, the medium was aspirated and cells were washed three times with PBS. Twelve milliliters of prewarmed basal medium consisting of DMEM/F12 containing 1x Primocin was added to each plate, and conditioned medium was collected after 24 h at 37°C and 5% CO2. Conditioned medium was centrifuged at 300 *xg* for 5 min at 4°C to remove floating cells and then at 2,000 *xg* for 10 min at 4°C to remove cellular debris. Halt phosphatase inhibitor (ThermoFisher 78420; 80 μL/sample) and protease inhibitor cocktail were added (Sigma P2714); 200 μL/sample, and samples were snap frozen in liquid nitrogen and shipped to Metware Bio for data-independent acquisition quantitative proteomics and analysis.

Briefly, samples were thawed on ice, supplemented with phenylmethylsulfonyl fluoride to a final concentration of 1 mM, centrifuged at 4,500 *xg* for 10 min at 4°C, and quantified using a bicinchoninic acid assay. Samples with low protein concentration were concentrated using 10-kDa molecular-weight-cutoff ultrafiltration devices. Thirty micrograms of protein per sample was denatured in 8 M urea, reduced with 5 mM dithiothreitol at 37°C for 45 min, and alkylated with 11 mM iodoacetamide for 15 min at room temperature in the dark. Samples were diluted with 25 mM ammonium bicarbonate and digested overnight at 37°C with trypsin and Lys-C. Peptides were acidified to pH 2– 3, desalted using C18 material, and quantified before LC-MS/MS analysis.

Peptides were separated using a NanoElute UHPLC system with an IonOpticks 25 cm x 75 μm C18 analytical column containing 1.6 μm particles. The column was maintained at 50°C, and 300 ng peptide was analyzed at a flow rate of 300 nL/min using a 47-min gradient. Mass spectrometry was performed on a Bruker timsTOF HT using diaPASEF acquisition.

Raw diaPASEF data were analyzed using DIA-NN version 1.8.1 with a library-free workflow. Spectra were searched against the UniProt human proteome database UP000005640 downloaded May 4, 2023. Deep-learning-based spectral prediction and match-between-runs were enabled, and precursor- and protein-level identifications were filtered at a 1% false-discovery rate. Proteins were quantified using the MaxLFQ algorithm based on Razor and unique peptides. Protein abundances were normalized within each sample relative to the median protein intensity.

For each pairwise group comparison, fold change was calculated from the mean normalized abundance across biological replicates, and p values were calculated using a t-test. Differentially abundant proteins were defined as having a fold change ≥1.5 or ≤0.6667 and p ≤0.05. A total of 7,623 peptides corresponding to 1,079 identified and quantified proteins were detected. For the analyses presented in the main text, proteins were subsequently restricted to those annotated as extracellular or plasma membrane associated.

### Loss of Function and Gain of Function Experiments

For gain of function experiments, myoblast lines were plated as described and incubated in differentiation medium in the presence of recombinant active CATHEPSIN L (R&D Systems, 10 nM) or holo-TRANSFERRIN (Millipore Sigma, 2.5 mg/mL). For inhibitor experiments, myoblast lines were incubated in differentiation medium with irreversible CATHEPSIN L inhibitor (Millipore Sigma, Z-FY(t-Bu)-DMK; CTSLi) or a neutralizing antibody raised against TRANSFERRIN (ThermoFisher, Clone HTF14, 10 μg/mL). After differentiation, cells were stained for MYOGENIN and MYOSIN HEAVY CHAIN and quantified as above. For myoblast-only loss-of-function experiments, CTSLi was used at a high dose (10 μM) to test whether CATHEPSIN L activity is required for myogenic differentiation under standard differentiation conditions. Because this concentration directly impaired myogenesis in the absence of conditioned media, a 100-fold lower concentration (100 nM) was used in FAP-conditioned media experiments to minimize direct inhibition of intrinsic myoblast differentiation and instead assess the contribution of CATHEPSIN L-dependent activity within conditioned media. Cells were stained for MYOGENIN and MYOSIN HEAVY CHAIN and analyzed as above.

### Large Language Model Use

This manuscript was edited by CHOP GPT v5.2.

### Statistical analysis of data

For all analyses other than mass spectrometry, statistical testing was performed using GraphPad Prism version 10.6.1. Comparisons among control, IC2 LOM, and pUPD11 groups were performed using one-way ANOVA. All tests were two-sided, and p<0.05 was considered statistically significant. Data are presented as mean ± SEM. Each data point in graphs represents one independent biological patient-derived FAP line or independent FAP-conditioned media-myoblast combination, as specified.

## Data Availability

The numerical source data underlying the graphs are provided in Supplementary Data 1. Raw mass-spectrometry data are available through the FaceBase repository under DOI 10.25550/AV-Z5Z6.

## Acknowledgements

We thank patients and families for contributing samples for this study. We thank Metware Bio for conducting mass spectrometry and subsequent data analysis/figure generation. We thank Rose Pradieu for specimen processing and Gracie Guerin, Sanya Dudeja, and Kirti Mhatre for assistance with data collection. We also thank the University of Pennsylvania Genetic Diagnostic Library and Dr. Arupa Ganguly for assistance with tongue sample methylation analysis. This work was supported by the Penn Center for Musculoskeletal Disorders pilot grant P30 (AR069619), CHOP Academic Enrichment Funding, the Berman Family Fund for Beckwith-Wiedemann Syndrome Research, the Cornerstone Beckwith-Wiedemann Syndrome Research Fund, the Victoria Fertitta Fund as part of the Lorenzo Turtle Sartini Jr. Endowed Chair in Beckwith-Wiedemann Syndrome Research, and NIH R01DE033646 to J.M.K.

## Author Contributions

E.D.T. and J.M.K. conceived the study. E.D.T., S.P., M.F., A.T.N., M.B. performed the methodology. E.D.T, S.P., M.F., A.T.N. performed the investigation. E.D.T. validated the data. E.D.T., S.P., M.N., M.F., A.T.N., G.K-S, M.B. performed the formal analysis. E.D.T. wrote the manuscript. All of the authors reviewed and edited the manuscript. J.M.K. acquired the funding. D.K., H.K., J.M.K. acquired the resources. J.M.K. supervised the study.

