## Supplementary for "Fibroadipogenic Progenitor Cells Contribute to Tongue Skeletal Muscle Hypertrophy in Beckwith-Wiedemann Syndrome"

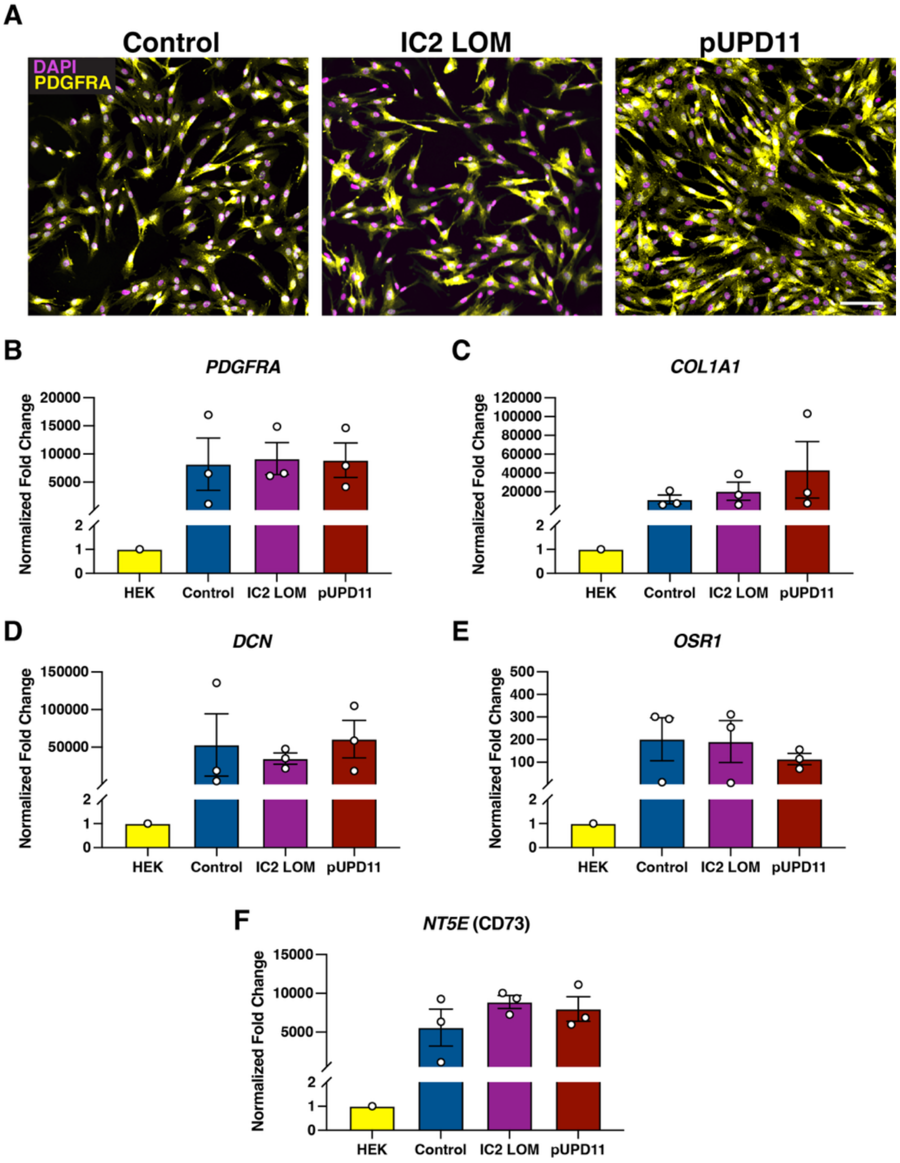


**Figure S1. Characterization of tongue-derived FAPs.**

**(A)** Representative images of tongue-derived FAPs following magnetic-activated cell sorting (MACS) for PDGFRA-positive cells. Isolated cells were plated in vitro and stained for PDGFRA to confirm enrichment of the FAP population. PDGFRA, yellow; nuclei, DAPI magenta. Scale bar: 100 µm. **(B-F)** Quantitative real-time PCR analysis of PDGFRA-positive cells isolated by MACS. Expression of the FAP markers *PDGFRA* (B), *COL1A1* (C), *DCN* (D), *OSR1* (E), and *NT5E* (F) was assessed to confirm population identity. Data were normalized to HEK cells. Each data point represents one primary FAP line derived from a unique patient.

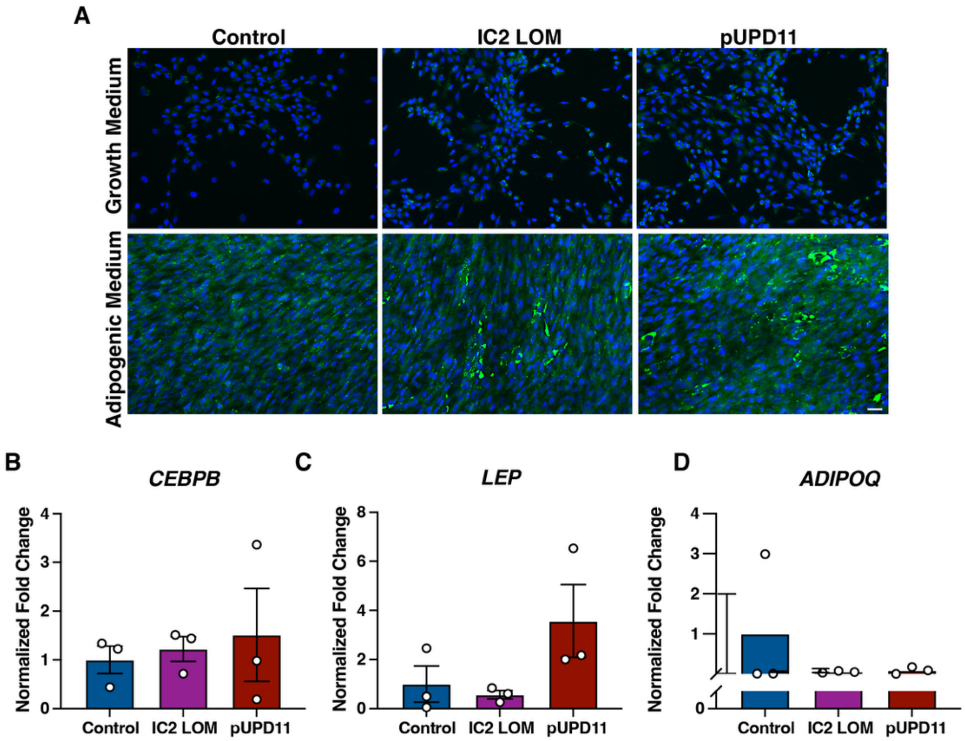


**Figure S2. Adipogenic differentiation of tongue-derived FAPs.**

**(A)** Representative images of control, IC2 LOM, and pUPD11 FAPs cultured in growth medium or adipogenic medium and stained with BODIPY 493 (green) to label neutral lipids and Hoechst (blue) to label nuclei. Scale bar: 50 μm. (B-D) Quantitative real-time PCR analysis of adipogenic gene expression, including *CEBPB* **(B)**, *LEP* **(C)**, and *ADIPOQ* **(D)**. Values were normalized to *UBC* expression.

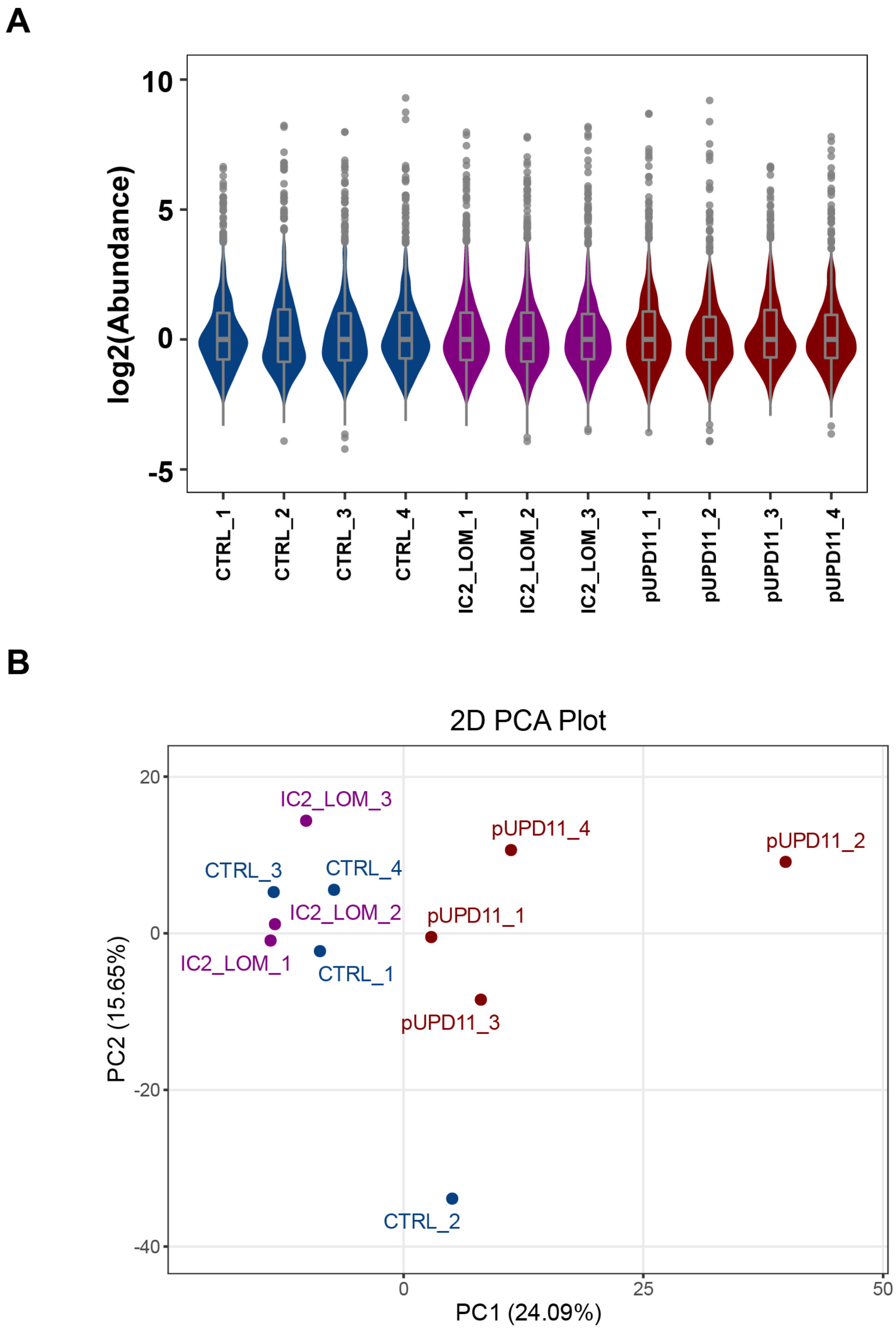


**Figure S3. Distribution and abundance of proteins identified in FAP secretomes.**

**(A)** Principal component analysis of secretomes from control (n = 4), IC2 LOM (n = 3), and pUPD11 (n = 4) samples. **(B)** Box plots showing protein abundance distributions across the secretome samples shown in (A).

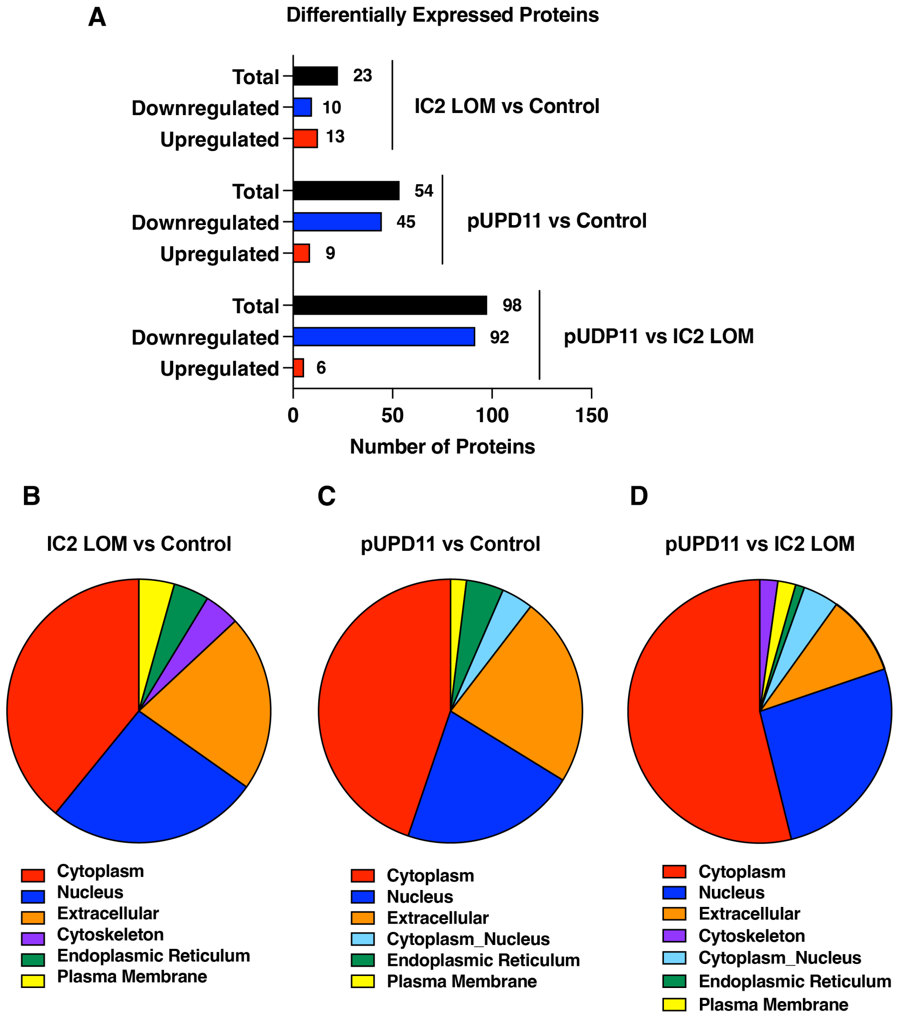


**Figure S4. Differentially expressed proteins in control, IC2 LOM, and pUPD11 FAP secretomes.**

**(A)** Quantification of upregulated and downregulated proteins across pairwise comparisons of experimental group secretomes. **(B)** Subcellular localization of differentially expressed proteins in IC2 LOM relative to control. **(C)** Subcellular localization of differentially expressed proteins in pUPD11 relative to control. **(D)** Subcellular localization of differentially expressed proteins in pUPD11 relative to IC2 LOM.

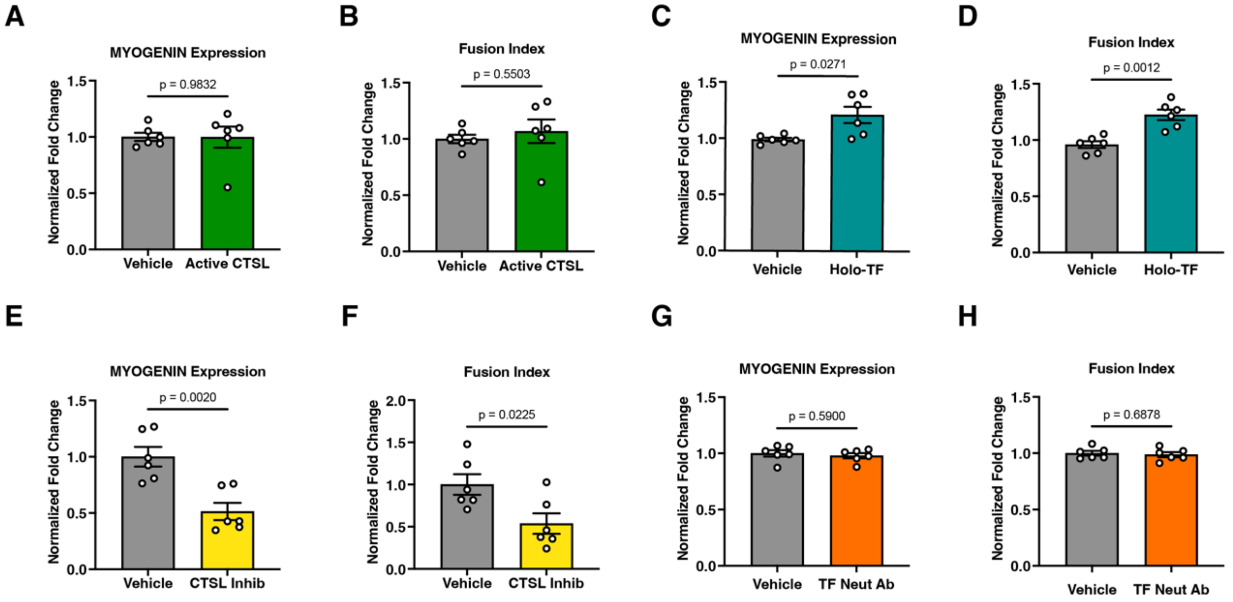


**Figure S5. Effects of CATHEPSIN L or TRANSFERRIN modulation on late stage myogenesis.**

**(A)** Quantification of MYOGENIN-positive nuclei in male and female human myoblast lines following exposure to recombinant active CATHEPSIN L. Each data point represents one biological replicate. **(B)** Quantification of fusion index in the same myoblast lines following exposure to recombinant active CATHEPSIN L. **(C,D)** Quantification of MYOGENIN-positive nuclei and fusion index, respectively, in male and female human myoblast lines following exposure to holo-TRANSFERRIN. **(E,F)** Quantification of MYOGENIN-positive nuclei and fusion index, respectively, in male and female human myoblast lines following exposure to an irreversible CATHEPSIN L inhibitor. **(G,H)** Quantification of MYOGENIN-positive nuclei and fusion index, respectively, in male and female human myoblast lines following exposure to a TRANSFERRIN-neutralizing antibody.

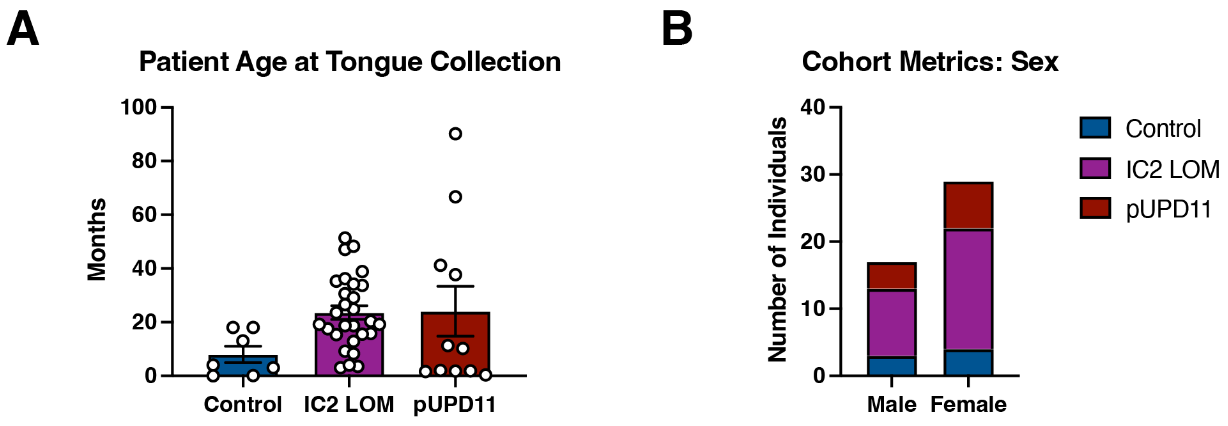


**Figure S6. Patient cohort characteristics for all samples in the study.**

(A) Patient age at the time of tongue specimen collection (for histology and FAP isolation).

(B) Patient sex for collected specimens (for histology and FAP isolation).

**Table S1: BWS Tongue Methylation Data FAP Isolation**

| Tongue sample ID | % methylation IC1 | % methylation IC2 | % mosaicism |
|---|---|---|---|
| IC2 LOM 1 | 50.95 | 12.97 | 74.06 |
| IC2 LOM 2 | 50.91 | 7.06 | 85.88 |
| IC2 LOM 3 | 51.17 | 6.57 | 86.86 |
| pUPD11 1 | 70.17 | 31.22 | 38.95 |
| pUPD11 2 | 78.73 | 22.61 | 56.12 |
| pUPD11 3 | 58.35 | 40.49 | 17.86 |
| pUPD11 4 | 84.13 | 19.42 | 64.71 |
